# Chromatin remodeling in peripheral blood cells reflects COVID-19 symptom severity

**DOI:** 10.1101/2020.12.04.412155

**Authors:** Nicholas S. Giroux, Shengli Ding, Micah T. McClain, Thomas W. Burke, Elizabeth Petzold, Hong A. Chung, Grecia R. Palomino, Ergang Wang, Rui Xi, Shree Bose, Tomer Rotstein, Bradly P. Nicholson, Tianyi Chen, Ricardo Henao, Gregory D. Sempowski, Thomas N. Denny, Emily R. Ko, Geoffrey S. Ginsburg, Bryan D. Kraft, Ephraim L. Tsalik, Christopher W. Woods, Xiling Shen

**Author notes:** These authors contributed equally to this work (Nicholas S. Giroux, Shengli Ding, Micah T. McClain). Correspondence to: (CWW), (XS).

## Abstract

SARS-CoV-2 infection triggers highly variable host responses and causes varying degrees of illness in humans. We sought to harness the peripheral blood mononuclear cell (PBMC) response over the course of illness to provide insight into COVID-19 physiology. We analyzed PBMCs from subjects with variable symptom severity at different stages of clinical illness before and after IgG seroconversion to SARS-CoV-2. Prior to seroconversion, PBMC transcriptomes did not distinguish symptom severity. In contrast, changes in chromatin accessibility were associated with symptom severity. Furthermore, single-cell analyses revealed evolution of the chromatin accessibility landscape and transcription factor motif occupancy for individual PBMC cell types. The most extensive remodeling occurred in CD14+ monocytes where sub-populations with distinct chromatin accessibility profiles were associated with disease severity. Our findings indicate that pre-seroconversion chromatin remodeling in certain innate immune populations is associated with divergence in symptom severity, and the identified transcription factors, regulatory elements, and downstream pathways provide potential prognostic markers for COVID-19 subjects.

**One sentence summary:** Chromatin accessibility in immune cells from COVID-19 subjects is remodeled prior to seroconversion to reflect disease severity.

## Introduction

Coronavirus disease 2019 (COVID-19), caused by severe acute respiratory syndrome coronavirus 2 (SARS-CoV-2) infection, manifests with highly variable symptom severity (*1*). Infected subjects demonstrate clinical trajectories that range from remaining asymptomatic to developing lifethreatening illness. The development of specific antibodies (IgG) against SARS-CoV-2 marks an inflection point in a COVID-19 patient’s disease progression, indicating a transition from innate immunity to acquired immunity (*2, 3*). IgG seroconversion typically occurs within two weeks of symptom onset and roughly coincides with the time that patients without critical illness will see clinical improvement (*4-6*). However, limited data are available examining the COVID-19 peripheral blood immune response in the context of seroconversion.

Several recent publications described bulk or single cell RNA-sequencing (scRNA-seq) to profile transcriptomic responses in immune cells of subjects with COVID-19 (*7-11*). Specifically, they reported suppressed immune responses in subjects with mild symptoms, as indicated by deficient expression of Type I and III interferons (*12*). Subjects with more severe disease demonstrated upregulation of pro-inflammatory factors, including IL-6 and TNF-alpha (*13*). The number of monocytes and associated IL-6, CCL2, and CCL8 production in the peripheral blood are also elevated in subjects with severe COVID-19 (*14, 15*). However, no investigation of the association between molecular profiles in PBMCs prior to seroconversion and symptom severity has been performed.

To test the hypothesis that the landscape of chromatin accessibility harbors biomarkers that define early molecular mechanisms underpinning divergent immunologic responses in SARS-CoV-2 infection, we performed bulk and single-cell RNA-seq and ATAC-seq on longitudinal PBMC samples from subjects before and after seroconversion.

## Results

PBMCs were collected from healthy controls, uninfected close contacts (CC), and COVID-19 subjects and profiled using bulk and single-cell RNA-seq and ATAC-seq sequencing (Fig. 1A). PBMCs from healthy subjects (n=7) were collected before the pandemic started. Infection with SARS-CoV-2 was confirmed using polymerase chain reaction (PCR) on nasopharyngeal (NP) swab samples, and serology testing for IgG against the SARS-CoV-2 spike domain was performed for each COVID-19 subject on study collection days. COVID-19 subjects included in this cohort were out-patients, and molecular profiling was performed at three timepoints, corresponding to early (IgG-), mid, and late (IgG+) acute disease within a 14-day window (Fig. 1B). At the mid timepoint, IgG seroconversion was observed in only some subjects; by the late timepoint, all subjects were observed to be IgG+ (Fig. 1B). CC subjects (n=7) were profiled at three timepoints and were PCR negative and IgG-at all timepoints. Subject symptoms were recorded for 39 symptom categories, and the total score was used to stratify the subjects into mild symptom (MS, n=7; mean score = 12.9) and pronounced symptom (PS, n=7; mean score = 33.6) cohorts. Complete subject demographics and clinical metadata are summarized in Tables S1-2.

**Figure 1:**
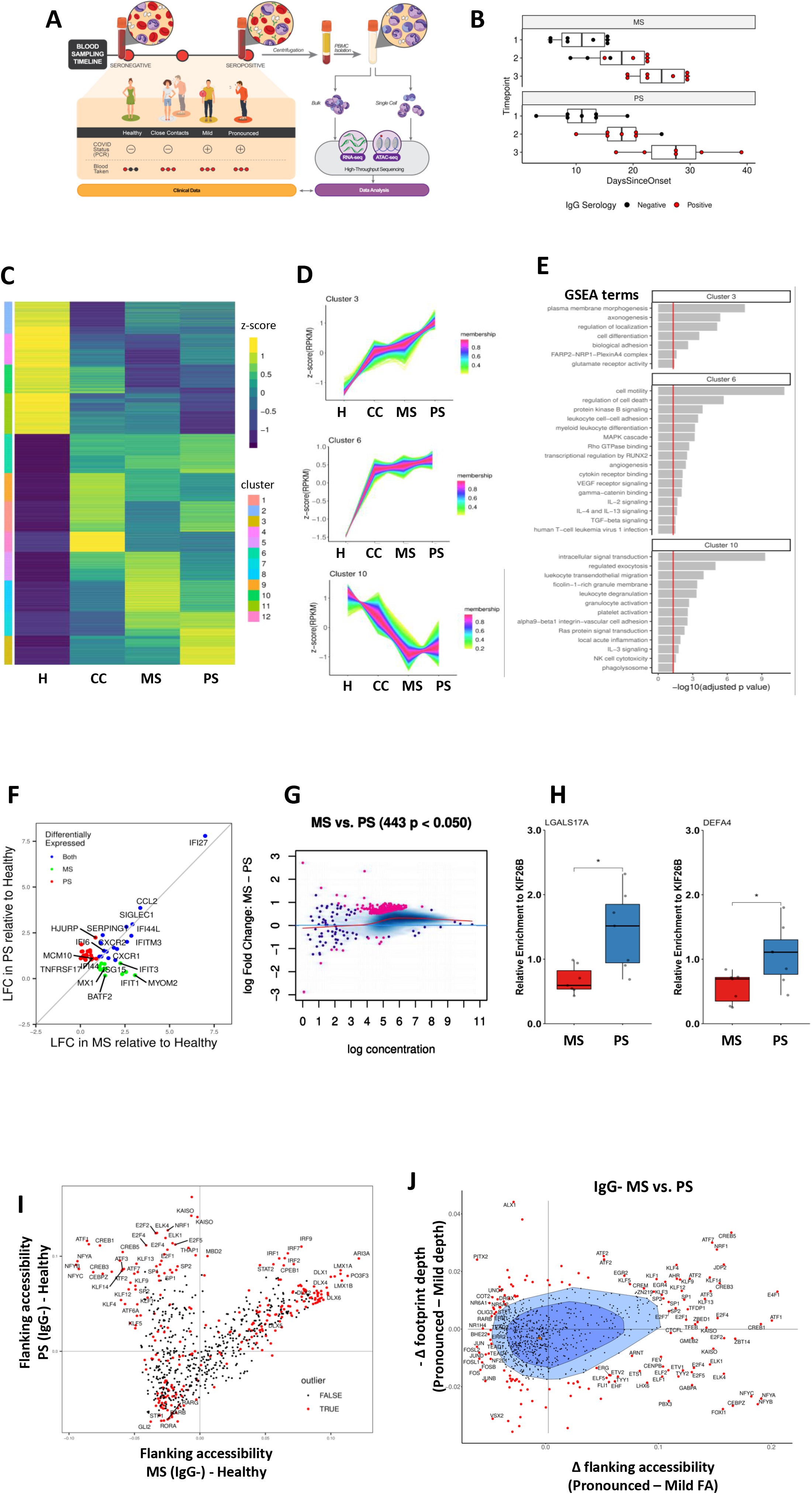
Chromatin accessibility in peripheral immune cells distinguishes COVID-19 symptom severity. **(A)** Summary of COVID-19 subject cohorts and multi-omic profiling experimental design. **(B)** Longitudinal timepoints available for COVID-19 subject cohort and IgG serological testing data. **(C)** Differential analysis of chromatin accessibility peaks from seropositive COVID-19 subjects identifies unique epigenetic biomarkers corresponding to symptom severity. Accessibility count data were normalized to reads per kilobase per million mapped reads and converted to z-scores for each group. **(D)** Differentially accessible peaks clustered by fuzzy c-means clustering with n=12. Membership indicates similarity of a peak to the centroid for each cluster. **(E)** Gene set enrichment analysis performed for peaks with membership > 50% for each cluster. Selected significant pathways plotted by -log(adj_p). **(F)** Differential gene expression of MS and PS subjects compared to a common healthy control baseline. **(G)** Accessible chromatin biomarkers that distinguish seronegative MS and PS subjects. Significant peaks have p < 0.05 and are plotted in pink. **(H)** ATAC-PCR enrichment of PS-specific markers LGALS17A and DEFA4 relative to internal control KIF26B. ATAC-seq libraries from seronegative MS (n=7) and PS (n=7) subjects were used as input. * p < 0.05 **(I)** Transcription factor footprinting analysis comparing motif flanking accessibility in MS and PS subjects compared to healthy controls. Outliers were determined by distance from BaGFoot center point. **(J)** BaGFoot comparing motif occupancies between MS and PS subjects. Elevated flanking accessibility and increased footprint depth indicates higher transcription factor occupancy at genome-wide motifs.

Samples from each COVID-19 (MS and PS) subject’s first IgG+ timepoint were compared with healthy controls and CC subjects to determine whether differential chromatin accessibility in PBMCs can stratify subjects by symptom severity (Fig. 1C). A total of 6168 significantly differentially accessible peaks were identified that distinguished each group in a pairwise comparison. Patterns of correlated chromatin peak accessibility emerged following analysis with soft clustering, indicating trends of increased accessibility that correspond with differential symptom severity (Fig. 1D, Fig. S1A). Clusters 3 and 6 were composed of peaks with decreased accessibility in healthy controls and increased accessibility following exposure (CC) or infection (MS and PS). Conversely, cluster 10 was composed of peaks with a negative correlation between accessibility and disease severity. This observed dependence on subject symptom severity suggested that the epigenome harbors biomarkers associated with COVID-19 severity. To evaluate whether the identified clusters contribute to disease variability, we performed gene set enrichment analysis (GSEA) for each cluster of peaks (Fig. 1E). Peaks with increased accessibility in MS and PS subjects were enriched in pathways related to interleukin signaling and regulation of cell differentiation and morphology. In contrast, healthy controls had the highest accessibility in peaks that were enriched for genes related to exocytosis, transendothelial migration, and vascular adhesion.

We compared bulk RNA-seq of PBMCs from the CC, MS, and PS cohorts at the earliest IgG-timepoint to the healthy controls. Activation of interferon response genes (IFI27, SIGLEC1, IFI44L) was observed in both seronegative MS and PS subjects compared to healthy controls (Fig. 1F, S1B) (*9*). However, these early transcriptomic differences do not reliably distinguish between MS and PS subjects (Fig. 1F).

Regulatory chromatin containing transcription factor motifs becomes accessible prior to downstream gene expression (*16*). To test whether chromatin changes can better distinguish disease severity, differential chromatin accessibility analysis was applied to the earliest seronegative samples from COVID-19 subjects compared to healthy controls. A set of 443 peaks of differentially accessible chromatin identified in seronegative subjects differentiates mild from more pronounced disease severity (Fig. 1G, S1C). As proof-of-principle, ATAC-PCR was performed to confirm enrichment of peaks annotated to LGALS17A and DEFA4 that were differentially accessible in PS subjects (Fig. 1H, S1D). Identification of an association between LGALS17A, a known interferon-stimulated gene, DEFA4, a marker of neutrophil activation in COVID-19 subjects, and disease severity confirmed the observations of the differential analysis (*17, 18*).

Motif occupancy analysis was applied to samples from the seronegative timepoint to identify regulatory transcription factors that bind newly accessible chromatin and poise gene expression. The seronegative timepoints from MS or PS subjects were compared to healthy controls using bivariate analysis to estimate the change in flanking accessibility and footprint depth at each motif (Fig. S1E-F). A group of transcription factors motifs with increased accessibility in PS subjects compared to MS subjects was identified by comparing samples from COVID-19 subjects before IgG seroconversion to a common baseline of healthy controls, in contrast to a second group of transcription factor motifs that demonstrated similar accessibility in both MS and PS subjects (Fig. 1I). The differential occupancies in the first group identified alternative regulation of myeloid activation pathways in MS vs. PS subjects. For example, KLF and CREB transcription factor families, which are known to regulate monocyte-macrophage polarization, have elevated motif occupancy in PS subjects compared to MS (*20, 21*). In the second group, CPEB1 displacement in both MS and PS subjects, marked by an increase in motif flanking accessibility and decrease in footprint depth, suggests involvement of the IL-6 and NFkB signaling pathways which was not observed in healthy controls (S1E-F) (*19*). Direct comparison of seronegative MS and PS subjects also demonstrated enrichment of KLF and CREB motif occupancy in PS subjects and served to further characterize differential regulation of immunity in each cohort (Fig. 1J). Furthermore, MS subjects were characterized by elevated AP-1 and C/EBP motif occupancies, which are known to negatively regulate IFN-gamma (*22, 23*).

The robust chromatin accessibility signature that was detected in the bulk ATAC-seq datasets confirms that PBMCs undergo extensive chromatin remodeling in response to SARS-CoV-2 infection. To understand how each cell type contributes to this signature and track the evolution of the chromatin landscape of each cell type during seroconversion, we performed single-cell ATAC-seq profiling of five subjects from each of the healthy, MS, and PS cohorts (Fig. 2A, Fig. S2A-F, Table S3). Cell type annotations were transferred from complementary single-cell gene expression libraries using canonical-correlation analysis, and expression of marker genes for each cell type was confirmed (Fig. S2G, Fig. S3A-E) (see Methods). Differential transcription factor motif occupancy analysis was applied to MS and PS PBMCs to identify shared regulatory mechanisms activated before seroconversion. Increased occupancy at transcription factor motifs including the proinflammatory C/EBP family and BACH1 marked the seronegative timepoint in both MS and PS subjects and further confirmed that the chromatin landscape is being remodeled prior to seroconversion (Fig. 2B) (*24, 25*). Differential accessibility in motifs of SMAD3, a known regulator of TGF-beta signaling, suggests a potential molecular driver that differentiates PS and MS subjects (*26*). Following seroconversion, both MS and PS subjects exhibit an increased accessibility at motifs of the transcription factor KDM2B, which is known to promote IL-6 production via chromatin remodeling (*27*).

**Figure 2:**
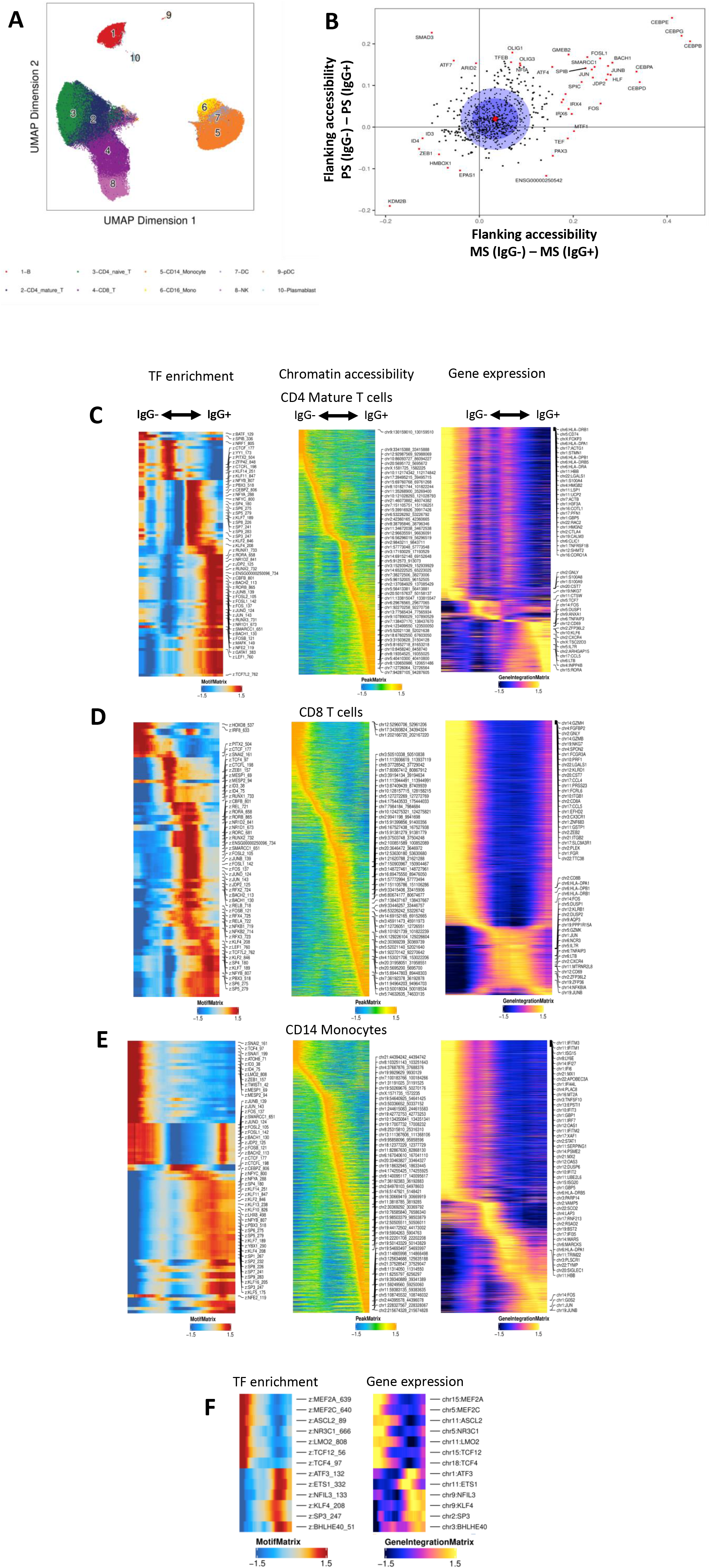
Epigenetic signature in cells collected from COVID-19 subjects evolves with disease progression. **(A)** UMAP plot of single-cell ATAC-seq datasets generated from healthy controls, uninfected CC, MS, and PS subjects. **(B)** BaGFoot analysis comparing transcription factor motif occupancies in PBMCs from MS and PS subjects collected prior to and after IgG seroconversion. (Red circle = average of all points; dark blue circle = 50% of all data; light blue circle = 75% of all data). Motifs with the top 5% change in flanking accessibility are plotted in red. **(C-E)** Supervised trajectory analysis using CD4+ mature T cells **(C)**, CD8+ T cells **(D)**, and CD14+ monocytes **(E)** collected from both MS and PS subjects. Differential transcription factor motif enrichment, chromatin accessibility, and gene expression were correlated with seroconversion. **(F)** Transcription factor regulators with correlated gene expression and transcription factor motif accessibility identified in CD14+ monocytes.

To identify the cell types with the most significant chromatin remodeling over time, we applied a global trajectory analysis to major cell types, including CD4+ mature T cells, CD8+ T cells, and CD14+ monocytes, represented in the scATAC-seq datasets (Fig. 2C-E). Similar analysis was also applied to CD4+ naïve T cells and NK cells, which contribute to the epigenetic signal to a lesser extent (Fig. S4A-B). Cells from MS and PS subjects were pooled into early (IgG-), mid, and late (IgG+) timepoints in pseudotime using a supervised trajectory algorithm (see Methods). Transcription factor motif enrichment, chromatin accessibility, and differential gene expression analyses were then performed for each individual cell population. A monotonic trend correlated with pseudotime was observed for each trajectory, demonstrating the progression of the host response in peripheral blood cells during development of acquired immunity. The shift in differential transcription factor motif and peak accessibility associated with pseudotime preceded changes in gene expression for all cells analyzed. We identified transcription factor activators, defined as genes with highly correlated expression and motif accessibility that act as robust multi-omic biomarkers that develop over time, in CD14+ monocytes (Fig. 2F). Gene activity scores, estimated using accessibility counts, were analyzed to identify chromatin biomarkers associated with seroconversion, which showed a clear transition in CD14+ monocytes (Fig. S4C).

Comparison of pooled PBMCs collected from MS and PS subjects identified specific peaksets enriched on the day of study enrollment (IgG-) versus a timepoint 2-4 weeks later (IgG+) (Fig. S5A, Fig. 3A top). A total of 5615 peaks were identified as uniquely accessible at one of the three longitudinal (early, mid, and late) timepoints (Fig. 3A bottom). A similar experimental approach was applied to each major cell type to detect those with extensive chromatin remodeling prior to seroconversion (Fig. 3B-C, S5B). The only cell types with enrichment of significant peaks at the early, IgG-timepoint were CD14+ monocytes (and, to a much lesser extent, CD4+ naïve T cells), which are the main contributor to the IgG-signature in the PBMCs shown in Fig. 3A-B. The remaining cell types, including CD4+ mature T cells, CD8+ T cells, and NK cells, were not characterized by peaks detectable prior to seroconversion. Transcription factor motifs enriched in peaks accessible prior to seroconversion had the highest occupancy in cells from the myeloid lineage, specifically CD14+ monocytes, dendritic cells, and CD16+ monocytes (Fig. 3D-E). In contrast, motifs enriched in peaks accessible after seroconversion lack elevated occupancy in monocytes, instead showing the highest activity in B cells and plasmablasts (Fig. S5C). The enrichment of accessible transcription factor motifs established early further supports the observation that CD14+ monocytes underwent the most chromatin remodeling among PBMCs prior to seroconversion.

**Figure 3:**
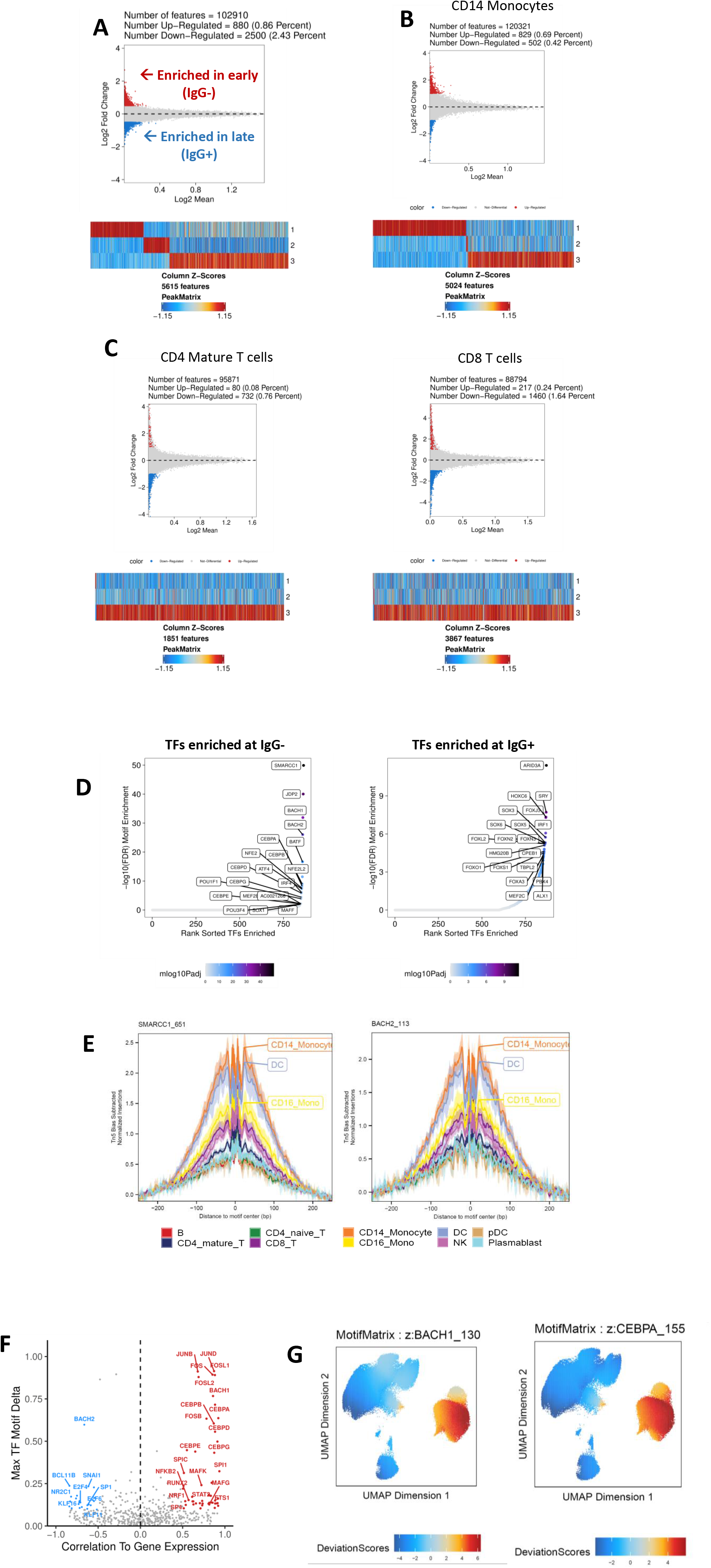
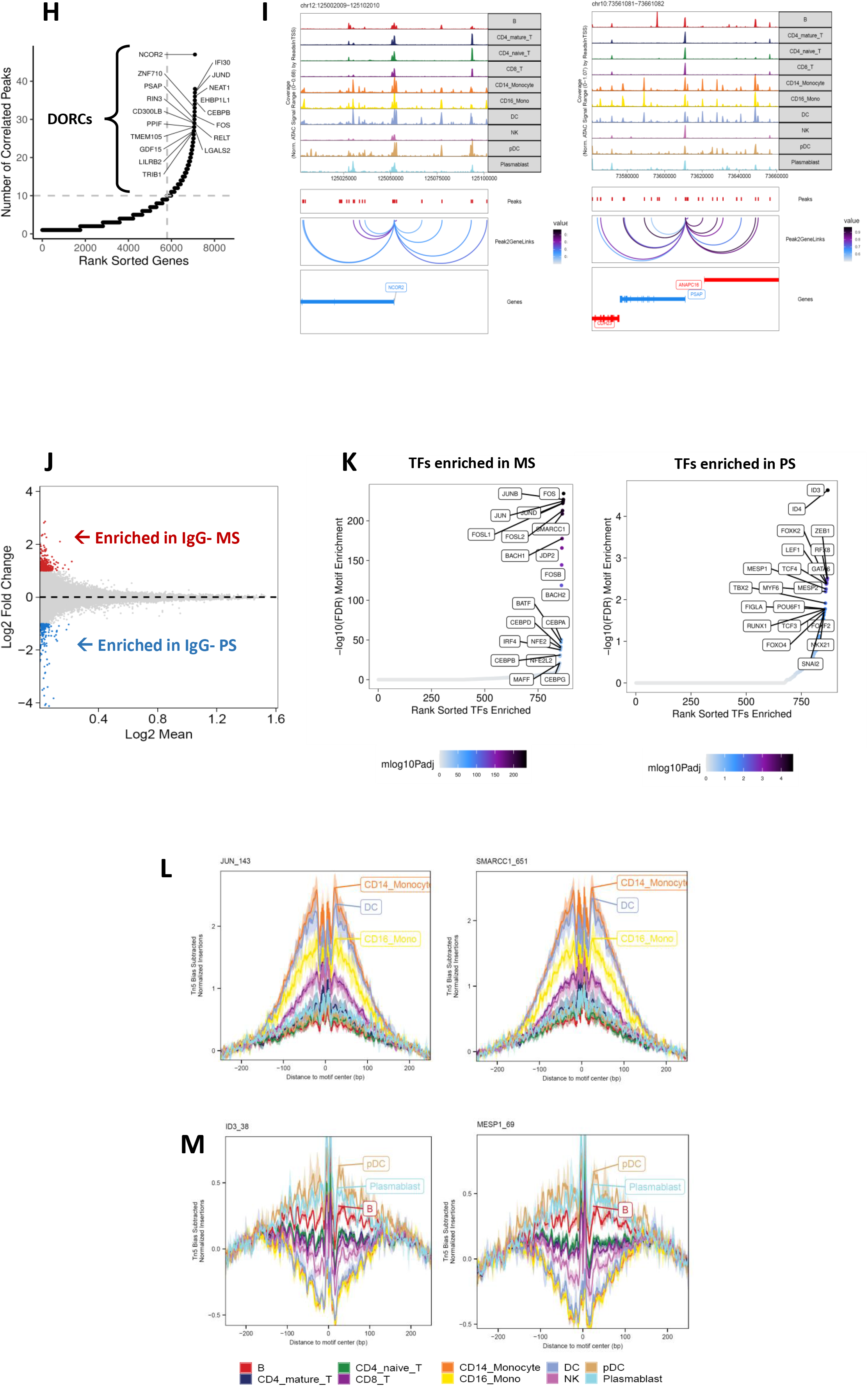
CD14+ monocytes undergo extensive chromatin remodeling prior to seroconversion and harbor severity-specific epigenetic biomarkers. **(A)** Top: Differentially accessible chromatin peaksets between early and late acute infection timepoints in all cells collected from both MS and PS subjects. Significance is defined as adjusted p < 0.05 and absolute LFC > 0.5. Bottom: Chromatin peaks with differential accessibility associated with time in all cells collected from MS and PS subjects. **(B)** Chromatin peaks with differential accessibility between early and late timepoints in CD14+ monocytes collected from MS and PS subjects. **(C)** Chromatin peaks with differential accessibility between early and late timepoints in CD4+ mature T cells and CD8+ T cells collected from MS and PS subjects. **(D)** Transcription factor motifs enriched in peaks specific to either early (IgG-) or late (IgG+) timepoints. **(E)** Transcription factors motifs enriched at the early timepoint have footprints with increased occupancy in CD14+ monocytes, dendritic cells, and CD16+ monocytes. **(F)** Transcription factor activators and repressors with correlated gene expression and motif accessibility associated with seroconversion in all cells. **(G)** Activator accessibility specifically enriched in CD14+ monocytes. Accessibility was overlaid on a UMAP of all cell types. **(H)** Peak-to-gene linkages computed for all cells collected from seronegative MS and PS subjects. Genes with > 10 linkages are defined as domains of regulatory chromatin (DORCs). Labeled genes are known to be regulated by a super-enhancer. **(I)** DORCs regulated by super-enhancers have increased accessibility in CD14+ monocytes. Peak-to-gene linkages are plotted with a correlation cutoff of 0.5. **(J)** Differentially accessible chromatin peaksets in all cells collected from seronegative MS and PS subjects. Significance is defined as adjusted p < 0.05 and absolute LFC > 1. **(K)** Transcription factor motifs enriched in peaks specific to either seronegative MS or PS subjects. **(L)** Seronegative MS-specific transcription factors have increased occupancy in CD14+ monocytes, dendritic cells, and CD16+ monocytes. **(M)** Seronegative PS-specific transcription factors have increased occupancy in pDCs, plasmablasts, and B cells.

Our analysis further identified transcription factors acting as activators or repressors, which have a positive or negative correlation between motif accessibility and gene expression, respectively, within the context of seroconversion (Fig. 3F). chromVAR deviation scores and accessibility at these activator motifs revealed an enrichment in CD14+ monocytes and other myeloid cells, including dendritic cells and CD16+ monocytes (Fig. 3F-G, Fig. S5D-E). A minority of activators (4 of 23) instead have enriched activity in B or T cells, potentially serving to poise these cells for future activation (Fig. S5D-E). These observations confirm that CD14+ monocytes are marked by a transcription factor activity signature that was established prior to seroconversion and undergo extensive chromatin remodeling during infection. Domains of regulatory chromatin (DORCs) were identified in all cell types collected from both seronegative MS and PS subjects by counting the number of peak-to-gene linkages for each gene in the scATAC-seq datasets (*16*). A total of 1109 genes with greater than 10 peak-to-gene linkages were defined as DORC genes, and those known to be regulated by a super-enhancer were labeled (Fig. 3H). These DORCs showed cell type-specific profiles and appeared to have higher accessibility than gene expression in these cells, suggesting that these genes are being primed for activation (Fig. S5F). Accessibility at the DORCs regulated by a super-enhancer was enriched in CD14+ monocytes and other myeloid cells, including dendritic cells and CD16+ monocytes (Fig. 3I, S5G).

We then determined the chromatin accessibility changes and corresponding cell types that discriminate seronegative MS and PS subjects. First, specific peaksets and transcription factor motifs enriched in all cells from seronegative MS and PS subjects were identified (Fig. 3J-K). MS-specific transcription factors, including subunits of AP-1, reproduce many of the same observations from bulk accessibility data comparing MS and PS subjects, and the motifs have the highest occupancy in CD14+ monocytes (Fig. 1I-J, Fig. 3K-L). In contrast, PS-specific transcription factor motifs have the highest occupancy in plasmacytoid dendritic cells (pDCs), plasmablasts, and B cells (Fig. 3K, 3M). These observations suggest that alternative regulation of PBMC chromatin by adaptive and innate immunity mechanisms is established prior to seroconversion and can be leveraged as potential biomarkers of disease severity.

CD14+ monocytes were identified as a key cell population that both responds to extensive chromatin reprogramming before seroconversion and distinguishes between subjects with mild or more pronounced symptoms. By applying a supervised trajectory algorithm and ordering monocytes from seronegative subjects in pseudotime from MS to PS, a strong correlation between differential chromatin accessibility and disease severity was identified (Fig. 4A). A differential gene expression and transcription factor motif enrichment signature distinguished CD14+ monocytes collected from MS and PS subjects (Fig. 4A). One timepoint was also profiled from six additional subjects who presented with critical symptoms (CS) and required intensive care unit (ICU) care (Table S4). Similar trends were observed in this independent cohort of IgG+, hospitalized subjects in which up-regulation of gene expression was observed in interferon response, IL-1 signaling, and neutrophil degranulation pathways (Fig. S6A-B). Additionally, we identified differential transcription factor motif enrichment and chromatin accessibility that distinguished CS subjects from MS and PS subjects (Fig. S6C).

**Figure 4:**
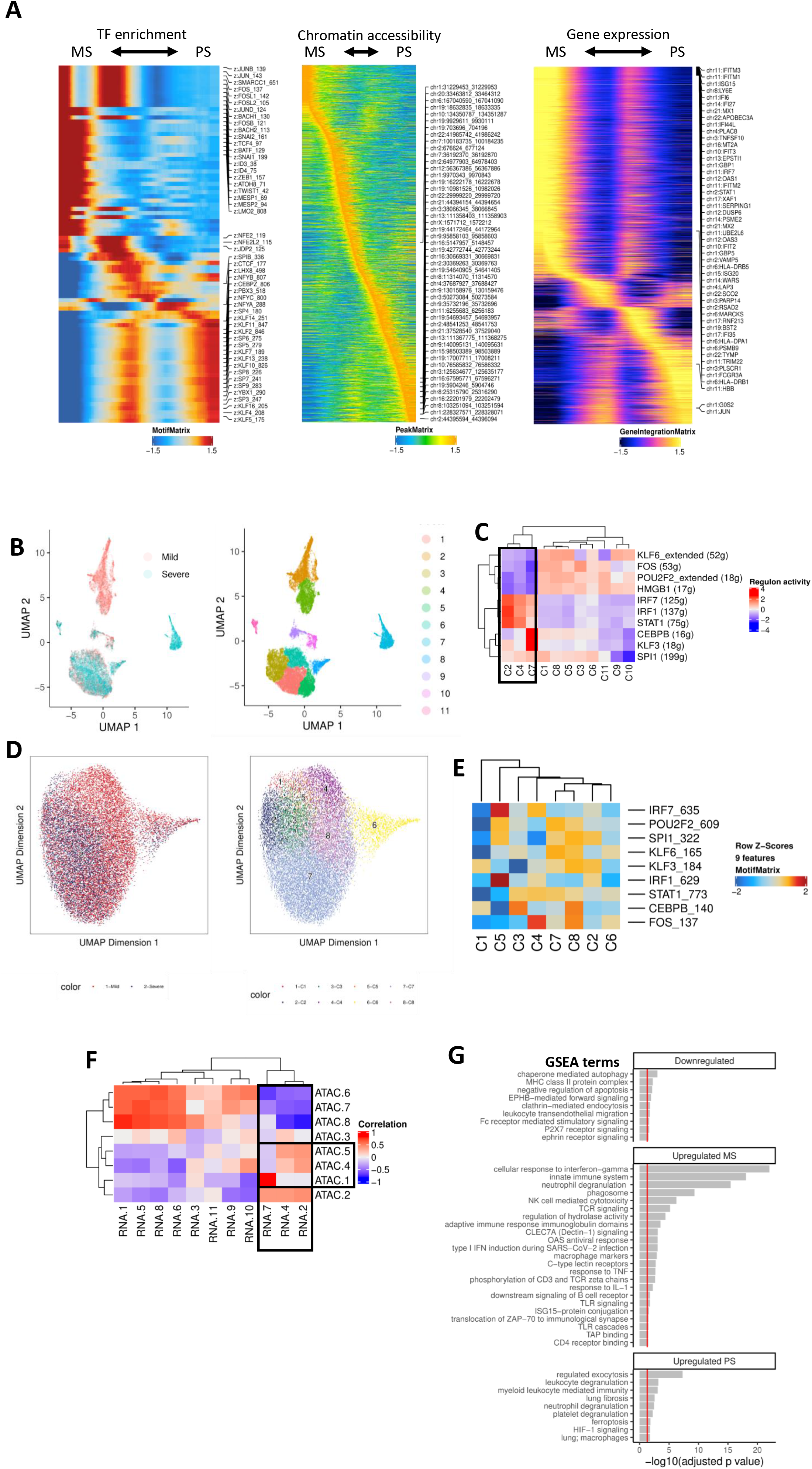
CD14+ monocytes distinguish seronegative MS and PS subjects. **(A)** Supervised trajectory analysis using CD14+ monocytes collected from seronegative MS and PS subjects. Differential transcription factor motif enrichment, chromatin accessibility, and gene expression were correlated with symptom severity. **(B)** scRNA-seq UMAP of all CD14+ monocytes collected from seronegative MS and PS subjects. UMAP colored by disease severity (left) and cluster number (right). **(C)** Regulon activity computed for each scRNA-seq cluster using SCENIC. Activity of the top 10 regulons is plotted as a heatmap for each cluster of cells from the UMAP. Black box indicates clusters of interest from UMAP in 4D. **(D)** scATAC-seq UMAP of all CD14+ monocytes collected from seronegative MS and PS subjects. UMAP colored by disease severity (left) and cluster number (right). **(E)** Transcription factor motif enrichment of regulons identified in 4G plotted for each scATAC-seq cluster. **(F)** Correlation plot using regulon activity to link clusters between scRNA-seq (columns) and scATAC-seq (rows). Black box indicates clusters of interest from UMAP in 4F. **(G)** Gene set enrichment analysis for regulons down-regulated (KLF6, FOS, POU2F2, HMGB1) or up-regulated (MS: IRF7, IRF1, STAT1; PS: CEBPB, KLF3) in scRNA-seq clusters 2, 4, and 7.

We next identified sub-populations of CD14+ monocytes characterized by both transcriptomic and epigenetic biomarkers that distinguish seronegative MS and PS subjects. Single-cell gene expression data for CD14+ monocytes were processed using the uniform manifold and projection (UMAP) algorithm and labeled by either subject disease severity or cluster (Fig. 4B). Clusters 2 and 4 identified a unique MS-specific sub-population of CD14+ monocytes from MS subjects, whereas cluster 7 identified a PS-specific sub-population. The gene regulatory networks were inferred for each scRNA-seq cluster to describe the activity of regulons, including transcription factors and their downstream target genes. Down-regulation of the KLF6, FOS, and POU2F2 regulons was observed in both the MS- and PS-specific monocyte subpopulations (Fig. 4C). The MS-specific sub-population was marked by up-regulation of the IRF7, IRF1, and STAT1 regulons, whereas the PS-specific sub-population was marked by up-regulation of the CEBPB and KLF3 regulons (Fig. 4C). Single-cell accessibility data for CD14+ monocytes was processed using a similar approach to identify clusters with correlated regulon activity (Fig. 4D, S6D). Activity at the transcription factor motifs was estimated, and the correlations between scRNA-seq and scATAC-seq clusters were calculated in a pairwise manner (Fig. 4E-F, S6E). Differential gene expression between the MS- and PS-specific sub-populations identified interferon-stimulated gene activation as a distinguishing feature of symptom severity (Fig. S6F-G). In addition to these transcriptomic markers, GSEA was then performed for the transcription factors with differential activity and their downstream target genes (Fig. 4G). Pathways identified using this epigenetic approach suggest alternative regulation of the adaptive immune response between MS and PS subjects. Differential regulation of pathways related to the transition from innate immunity, marked by the implication of dectin-1 and the toll-like receptor cascade, to adaptive immunity, suggested by activation of T and B cell receptor signaling, were observed in these sub-populations (*28-30*). These observations confirm that chromatin accessibility remodeling corresponds to symptom severity, as first indicated by differential transcription factor activity.

## Discussion

In this study, we identified an association between the evolution of the chromatin landscape in PBMCs and the severity of COVID-19 in a longitudinal cohort of subjects that underwent IgG seroconversion. We present evidence here that the epigenomes of PBMCs are remodeled extensively following infection by SARS-CoV-2 and reflect disease severity. Specifically, differential activity of transcription factors and chromatin accessibility prior to seroconversion were observed to distinguish disease severity, preceding the later transcriptional response. Upon analysis of each major cell type, we showed that CD14+ monocytes underwent extensive chromatin remodeling over time and harbor epigenetic biomarkers in seronegative subjects that distinguish disease severity. We further identified enrichment of DORC genes regulated by superenhancers in CD14+ monocytes which play a role in priming active chromatin states and cell fate decisions and observed sub-populations of CD14+ monocytes with severity-specific transcription factor activities (Fig. S6H) (*16*). Similar analysis applied to tissue-resident immune cell populations may further elucidate the factors driving divergence of disease severity (*31, 32*). These observations will inform new hypotheses to characterize the immunological host response to SARS-CoV-2 infection and be used to assess new strategies for treatment.

We demonstrated proof-of-principle that an ATAC-PCR-based assay can potentially be used as an epigenetic test to stratify disease severity (*33*). Development of a host-response assay that leverages the highly sensitive epigenetic biomarkers established early during infection and capable of predicting disease severity has the potential to fill an unmet clinical need in the care of COVID-19 patients (*34, 35*). It will be important to apply this analysis to a longitudinal cohort of subjects who go on to develop worsening symptoms and require critical care in order to generate data for a prognostic epigenetic assay.

In summary, we propose that evolution of the chromatin landscape in the peripheral blood of COVID-19 subjects primes the immunological host response mediated by CD14+ monocytes and drives the observed divergence in disease severity. We also demonstrate that these changes precede transcription of pathways related to adaptive immunity, and these observations can potentially be used to inform clinical approaches to COVID-19 patient treatment.

## Acknowledgments

We thank Drs. Eric van Gieson and Thomas Thomou for their advice throughout the project. We also thank Drs. Andrew Macintyer and Thomas Oguin, and their research teams for running, for performing COVID IgG serology and PCR viral load testing in the Core Services units of the Duke Regional Biocontainment Laboratory. We thank Raul Louzuo, Christina Nix, Tyffany Locklear, Pam Isner, Anna Mazur, Jack Anderson, Maria Miggs, Aleah Bowie, Julie Steinbrink, Robert Rolfe, and Jorge Prado for their contributions.

## Author contributions

NSG, SD, CWW, and XS designed the experiments. NSG performed the analysis of the sequencing data. NSG and XS interpreted the data and wrote the manuscript, which was critically revised by all remaining authors. MTM, TWB, and EP procured clinical samples and patient data. NSG, SD, HAC, GRP, EW, RX, and TR performed the RNA-seq and ATAC-seq profiling and ATAC-PCR assays. SB drafted the schematic figures. TC and RH provided advice for the analysis of the sequencing data. GDS and TND coordinated COVID subject sample processing, IgG serology and PCR assessments used to stratify subjects. BPN, ERK, GSG, BDK, ELT, and CWW contributed to clinical study design and collection.

## Data and materials availability

The sequencing datasets and related clinical metadata tables will be made available via the Gene Expression Omnibus (GEO) repository prior to publication.

## Supplementary Materials

Materials and Methods

Tables S1-S4

Figures S1-S6

## Materials and Methods

### IRB Approvals

Research protocols were approved by the relevant institutional IRBs and were performed in accordance with the Declaration of Helsinki. Written informed consent was obtained from all research participants or their legally authorized representatives (LAR).

### Subjects and PBMC sample collection

Subjects with confirmed or suspected SARS-CoV-2 infection, or their close contacts, were identified either in the hospital (Duke University Medical Center, Durham, NC; Duke Regional Hospital, Durham, NC; or Durham VA Health Care System, Durham, NC) or outpatient setting, and enrolled into the Molecular and Epidemiological Study of Suspected Infection protocol (MESSI, IRB Pro00100241). Where possible, longitudinal samples and data were collected through convalescence. The CC group consisted of seven participants with a close contact to someone in the same household with confirmed COVID-19 disease, but CC subjects had negative SARS-CoV-2 PCR and serological testing for at least two months after exposure.

Eight subjects with mild symptoms (MS) had a total symptom score <20 on the day of enrollment (mean = 12.8). Seven subjects with more pronounced symptoms (PS) had a total symptom score >20 on the day of enrollment (mean = 33.6). Six subjects were hospitalized due to COVID-19 and were categorized as having critical symptoms (CS) including three with acute respiratory distress syndrome (ARDS) requiring intensive care unit (ICU) care (*36*). All MS and PS COVID-19 subjects were longitudinally sampled from enrollment to convalescent phase. Seven healthy controls were asymptomatic and enrolled prior to the COVID-19 pandemic.

### SARS-CoV-2 IgG ELISA

Antibody response testing was performed using the anti-SARS-CoV-2 IgG ELISA assay (EUROIMMUN Medizinische Labordiagnostika AG, Lübeck, Germany) according to manufacturer’s instructions. Test results were evaluated by calculating the ratio of the OD (optical density) of the test sample over the OD of the calibrator sample. Ratio of <0.8 was interpreted as negative and ratio of 1.1 or greater as positive (ratio of 0.8 to <1.1 as indeterminate and not utilized in phenotyping).

### SARS-CoV-2 quantification by qRT-PCR

Nasal swab Viral Transport Medium (VTM) was aliquoted and cryopreserved from study subjects to determine SARS-CoV-2 N1 gene copy number by RT-PCR to stratify subjects as COVID PCR positive or negative. Viral RNA was extracted from 140 uL of VTM according to manufacturer’s instructions (QiaAmp Viral RNA minikit). SARS-CoV-2 nucleocapsid (N1) and human RNase P (RPP30) RNA copies were determined using 5 μL of isolated RNA in the CDC-designed kit (CDC-006-00019, Revision: 03, Integrated DNA Technologies 2019-nCoV kit). Standard quantitative RT-PCR (TaqPath 1-step RT qPCR Master Mix, Thermofisher) was run with test RNA and gene-specific standard curves (2e5 copy/mL – 20 copy/mL). Regression analysis was used to determine gene copy number and corrected to report copies/mL of VTM. Samples with a Ct value less than 35 are called as COVID PCR Negative and samples greater than or equal to 35 are called COVID PCR Positive.

### Collection of PBMCs

PBMCs were prepared using the Ficoll-Hypaque density gradient method. Whole blood was collected in ACD vacutainer tubes and processed within 8 hours. Blood was diluted 1:2 in PBS, layered onto the Ficoll-Hypaque in 50 ml conical tubes, and centrifuged at 420 g for 25 minutes. Buffy coat was collected and washed with D-PBS by centrifugation at 400 g for 10 minutes. Cell pellets were resuspended in D-PBS and washed again. PBMCs were assessed for viability and cell count using a Vi-Cell automated cell counter (Beckman-Coulter). PBMCs were adjusted to 10×10^6^ cells/ml in cryopreservation media (90% FBS, 10% DMSO) and then aliquoted into cryopreservation vials on ice. Cells underwent controlled freezing at −80C using CoolCell LX (BioCision) for 12-24 hours and were then transferred to liquid nitrogen vapor phase.

### RNA extraction, total RNA-seq, and analysis

For each sample, RNA was extracted from 300K cells using Zymo Direct-zol miniprep kit (Cat# R2051). RNA quality was assessed using Agilent DNA tape screen assay. The RNA Integrity Number (RIN) scores for all samples were > 7.0. Total RNA libraries were generated using NuGEN Ovation^®^ SoLo RNA-Seq Library Preparation Kit (Cat# 0500-96).

#### FASTQ processing

FASTQ files were generated from the NovaSeq BCL outputs and quality was assessed with FASTQC(*37*). Eukaryotic rRNA sequences were removed using SortMeRNA, and the remaining reads were aligned against the hg19 human reference genome using STAR and the following commands: STAR –genomeDir /path/to/STARIndex/ --sjdbGTFfile /path/to/gene.gtf –readFilesIn /path/to/R1.fastq /path/toR2.fastq –runThreadN 8 –twopassMode Basic –outWigType bedGraph – outSAMtype BAM SortedByCoordinate –readFilesCommand zcat –outReadsUnmapped Fastx – outFileNamePrefix $sampleID. Following alignment, the gene count matrix was generated using featureCounts (*38, 39*).

#### Differential gene expression analysis

Differentially expressed genes were identified between subjects with different disease severity using the ‘wald’ test in DESeq2(*40*). Subject sex and RNA-seq library batch were added as variables to the design formula to account for expected technical variation in the counts. Ribosomal protein genes and genes not annotated as protein coding in Ensembl were filtered from the gene count matrix. False discovery rate adjustment was performed for the p-values using the ‘BH’ method, and log2 fold change shrinkage was performed using the ‘ashr’ method. A gene was defined as significantly differentially expressed if the adjusted p-value < 0.05 and the shrunken absolute log2 fold changes > 1. Results from DESeq2 were passed to the EnhancedVolcano package to generate volcano plots with the same p-value and log2 fold change thresholds(*47*).

### ATAC-seq and analysis

Nuclei were extracted from frozen PBMCs. Briefly, 100K cells were spun down at 300 g for 5 minutes at 4°C. The supernatant was removed, and cells were mixed with 100 ul of lysis buffer (10mM NaCl, 3mM MgCl2, 10mM Tris-HCl pH7.4, 0.1% Tween-20, 0.1% Nonidet™ P40) and lysed on ice for 4 minutes. Wash buffer (1 mL; 10mM NaCl, 3mM MgCl2, 10mM Tris-HCl pH7.4, 0.1% Tween20) was added before nuclei spin at 500 g for 5 minutes at 4°C. ATAC-seq libraries were generated as presented earlier (*42*). Briefly, transposition mix (25 μl 2× TD buffer, 2.5 μl transposase (Tn5, 100 nM final), 22.5 μl water) (Illumina Cat# 20031198) was added to the nuclear pellet. The reaction was incubated at 37 °C for 30 minutes. Samples were purified using Qiagen MinElute PCR Purification Kit (Qiagen Cat#28004). DNA fragments were PCR amplified for a total of 10-11 cycles. The resulting libraries were purified again using Qiagen MinElute PCR Purification Kit. The libraries were sequenced with an Illumina Novaseq 6000 S4 flow cell using 100 bp paired-end reads.

#### FASTQ processing

FASTQ files were generated from the NovaSeq BCL outputs and used as input to the ENCODE ATAC-seq pipeline (https://github.com/ENCODE-DCC/atac-seq-pipeline) using the MACS2 peak-caller with all default parameters. Output narrowPeak files and aligned BAM files were used for downstream analysis.

#### Differential chromatin accessibility analysis

Differential accessibility was calculated between groups of subjects with different disease severity using the TCseq package(*43*). A peak count matrix was generated using a set of consensus peaks and the following parameter values: filter.type = “raw”, filter.value = 30, samplePassfilter = 6. Peaks that were differentially expressed (p < 0.05) in seropositive subjects were used as input to the comparison of healthy controls, CC, MS, and PS subjects. Differentially accessible peaks were defined as having a false discovery rate adjusted p-value < 0.1 and absolute fold change > 0.5. Clustering was applied to the differentially accessible peaks using the ‘cm’ method and k = 12. Peaks with a cluster membership > 0.5 were annotated to the closest gene using the ChIPseeker package and UCSC hg19 annotation database (available through the TxDb.Hsapiens.UCSC.hg19.knownGene package) (*44, 45*). Protein-protein interactions were estimated using STRING and the network was trimmed using k-means clustering to remove genes with few interactions (*46*). Functional profiling was performed for these genes using g:Profiler (*47*).

Outputs from the ENCODE pipeline were also used for analysis with the DiffBind package to estimate differential accessibility between seronegative MS and PS subjects (*48*). A consensus peakset was generated requiring minOverlap = 0.9 in all samples. Peak count normalization was applied using the following parameter values: background = TRUE, method = DBA_EDGER, normalize = DBA_NORM_LIB, library = DBA_LIBSIZE_FULL, offsets = TRUE. Significant differentially accessible peaks were defined as having p<0.05.

#### Bivariate genomic footprinting analysis

Bivariate genomic footprinting analysis was performed using the bagfoot package and all default parameter values (*49*). NarrowPeak files for all samples were merged to generate a consensus peakset for motif enrichment analysis. BAM alignment files were merged using picard MergeSamFiles and indexed with samtools (http://broadinstitute.github.io/picard/).

### scRNA-seq and analysis

Frozen PBMCs were thawed, and count and cell viability were measured by Countess II. The cell viability exceeded 80% for all samples except PBMC samples from CS subjects, which had viability between 70-80%. For single cell RNA-seq, 200K cells were aliquoted, spun down, resuspended in 30 ul PBS+0.04%BSA+0.2U/ul RNase inhibitor, and counted using Countess II. GEM generation, post GEMRT cleanup, cDNA amplification, and library construction were performed following 10X Genomics Single Cell 5’ v1 chemistry. Quality was assessed using Agilent DNA tape screen assay. Libraries were then pooled and sequenced using Illumina NovaSeq platform with the goal of reaching saturation or 20,000 unique reads per cell on average. Sequencing data were used as input to the 10x Genomics Cell Ranger pipeline to demultiplex BCL files, generate FASTQs, and generate feature counts for each library.

#### Dimensionality reduction and cell type annotation

Gene-barcode matrices generated using CellRanger count were analyzed using Seurat 3 with the default parameters unless otherwise specified (*50*). Cells with > 5% of reads mapping to the mitochondrial genome or > 2500 genes detected were removed from the analysis. Counts were log-normalized, and the top 2000 variable features were identified. Principal component analysis was performed using these variable genes, and the top 20 principal components were used for downstream analysis. UMAP dimensionality reduction was performed using the top 20 principal components identified using the Harmony package (*57*). Graph-based clustering was performed with resolution = 0.5. Cell types were inferred by using the DatabaseImmuneCellExpressionData() method from the SingleR package (*52*). Labels were confirmed by identification of differentially expressed genes using the FindAllMarkers() method.

#### Regulatory network inference

The scRNA-seq Seurat object was converted into a SingleCellExperiment and used as input to analysis with the SCENIC package (*53*). Cells from seronegative MS and PS subjects were reclustered using Monocle 3, and the top 100 marker genes were computed for each cell partition (*54*). The standard workflow for running the SCENIC analysis was then performed using the count matrix for these marker genes as input (https://github.com/aertslab/SCENIC). Briefly, GENIE3 was used to identify regulons of transcription factors and their downstream regulatory targets with correlated co-expression, and AUCell was then used to score the activity of these regulons in each cluster. The ‘top10perTarget’ co-expression parameter value was used to prune the list of scored regulons.

### scATAC-seq and analysis

PBMCs were thawed and aliquoted as mentioned above. Nuclei were extracted as for ATAC-seq. The single-cell suspensions of scATAC-seq samples were converted to barcoded scATAC-seq libraries using the Chromium Single Cell 5’ Library, Gel Bead and Multiplex Kit, and Chip Kit (10x Genomics). The Chromium Single Cell 5’ v2 Reagent (10x Genomics, 120237) kit was used to prepare single-cell ATAC libraries according to the manufacturer’s instructions. Quality was assessed using Agilent DNA tape screen assay. Libraries were then pooled and sequenced using Illumina NovaSeq platform with the goal of reaching saturation or 25,000 unique reads per nuclei on average. Sequencing data were used as input to the 10x Genomics Cell Ranger ATAC pipeline to demultiplex BCL files, generate FASTQs, and generate feature counts for each library.

#### scRNA-seq and scATAC-seq integration

Fragment file outputs generated using CellRanger ATAC count were analyzed using ArchR following the standard workflow and with default parameters unless otherwise specified (*55*). Cells with a transcription start site enrichment score < 4, cells with fewer than 1000 detected fragments, and putative doublets were removed from downstream analysis. Dimensionality reduction was computed using iterative latent semantic indexing (LSI), and batch effect correction was applied using Harmony. Graph-based clustering was performed using the FindClusters() method from Seurat 3 with resolution = 0.8. UMAP embeddings were calculated with the top 30 principal components from either LSI or Harmony. Constrained integration was performed using the addGeneIntegrationMatrix() method and scRNA-seq cell type annotations were used to label the identify of scATAC-seq clusters.

#### Feature and motif enrichment analysis

Peak calling was performed using MACS2 via the addReproduciblePeakSet() method in ArchR which uses pseudo-bulk replicates of cells grouped on a specific design variable. Differentially accessible peaks were identified between two groups and visualized using the ArchR methods getMarkerFeatures() and markerPlot(), respectively. Significance was defined as FDR <= 0.1 and absolute log2 fold change >= 0.5 unless otherwise specified. The ‘cisbp’ motif set was imported from TFBSTools using the ArchR addMotifAnnotations() method, and motif enrichment in differentially accessible peaks was performed using the peakAnnoEnrichment() method. Additionally, chromVAR deviation scores for these motifs were computed using the ArchR implementation (*56*). Motif footprinting was performed by measuring Tn5 insertions in genomewide motifs and normalized by substracting the Tn5 bias from the footprinting signal.

#### Integrative analysis with scRNA-seq

The correlations between chromVAR transcription factor deviation scores and gene expression data were calculated using the ArchR method correlateMatrices() to identify activators and repressors.

Peak-to-gene linkages were calculated using the addPeak2GeneLinks() method in ArchR using a correlation cutoff of 0.5 and resolution = 1. This approach uses low-overlapping cell aggregates to reduce noise that arises from doing correlative analyses with sparse scATAC-seq datasets. Peak-to-gene linkages were plotted against peak accessibility at DORC genes for each cell type. DORC genes were defined as gene loci with > 10 peak-to-gene linkages, and these sites were used as input to the web tool Seanalysis to identify regulation by a known super-enhancer in peripheral blood cells (*16, 57*).

The activity of the top-ranked transcription factor regulators that were correlated with scRNA-seq clusters was estimated for each scATAC-seq cluster. These activities were used to calculate Pearson correlation coefficients between scATAC-seq clusters (C1-C8) and scRNA-seq clusters (C1-C11) were calculated to identify scATAC-seq clusters with similar regulatory network activity.

#### Supervised pseudotime trajectory analysis

Cellular trajectories were established in a low-dimensional space using LSI embeddings and a user-defined trajectory backbone. For this study, a rough ordering of 2-3 groups of cells specified with a design variable was provided to the ArchR method addTrajectory(). Then, a k-nearest neighbors algorithm was used to order cells based on the Euclidean distance of each cell to the nearest cluster’s centroid. Cells were then assigned pseudotime value estimates, and a heatmap was plotted using differential feature z-scores that were associated with the pseudotime trajectory.

### ATAC-qPCR

The optimal primer regions for peaks of interests were designed by the ATACPrimerTool (APT) (https://github.com/ChangLab/ATACPrimerTool). The identification of the optimal primer regions and primer design were performed as previously described (*33*). In short, original .bam files and .bed files containing peak of interests were provided as inputs for the APT, and hg19 was used as the reference genome. Primers were designed by Primer3 Plus based on the APT identified optimal primer regions with the parameters listed in the original APT paper. The ATAC-qPCR was done by assembling a qPCR reaction containing 1ng ATAC-seq library, 125nM of forward and reverse primers and SYBR Green Master Mix with the following cycling conditions: 2 minutes at 98C, 40 cycles of 10 seconds at 98C, 20 seconds at 60C and 30 seconds at 72C. The two Universal Normalization Primers AK5 and KIF26B were used as internal control for the qPCR experiment.

Primers used:

Universal Normalization primers:

AK5 Forward primer: AGCGCGGAGACCACAG

AK5 Reverse Primer: CGGTGCAGCCCTCTTTC

KIF26B Forward primer: AAGCTCGGTGAAGGAGACAA

KIF26B Reverse primer: ACGAGGAAAGCGAGGGATAC

PS-enriched gene primers:

DEFA4 Forward primer: CACTGGGCTATGGAGGACTG

DEFA4 Reverse primer: AGCCGACTTCACTGCTCTG

LGALS17A Forward primer: GTGTGTGCTGGGATGTGACT

LGALS17A Reverse primer: CTGCTGTGTTGGGAGGAAAC

**Supplemental Table 1.** Subject serology and PCR results on days when PBMCs were collected for bulk assays. Summary of demographics for enrolled healthy controls, CC, MS, and PS subjects. Maximum severity score corresponds to the highest severity score during infection calculated from a survey of 39 symptom categories. (SE = standard error)

| Characteristics | Healthy controls | Close contacts (CC) | Disease Severity |  |
| --- | --- | --- | --- | --- |
|  |  |  | Mild (MS) | Pronounced (PS) |
| **Age**<br>Mean (Range), yr | 41.0 (27-60) | 42.9 (17-60) | 33.0 (26-60) | 34.0 (20-51) |
| **Gender**<br>(Male, %) | 57.1% | 57.1% | 62.5% | 42.9% |
| **Max severity score ± SE** | N/A | 9.9±2.8 | 12.9±2.1 | 33.6±2.4 |

**Supplemental Table 2.** Subject serology and PCR results on days when PBMCs were collected for bulk assays. Serology testing was performed for IgG against the SARS-CoV-2 spike protein on each day of PBMC collection. Collection days are relative to study enrollment with sequencing profiling and specified for each subject.

| Disease Severity | Alias ID | Race | Ethnicity | Days of PBMC collection | Serology Test (IgG) | PCR |
| --- | --- | --- | --- | --- | --- | --- |
| **Healthy** | 22CBD6 | White | NR* | N/A | N/A | N/A |
|  | 77DA77 | White | NR | N/A | N/A | N/A |
|  | 4F2EAD | White | NR | N/A | N/A | N/A |
|  | 1FD3AC | White | NR | N/A | N/A | N/A |
|  | F8FF1A | White | NR | N/A | N/A | N/A |
|  | 56E219 | Black | NR | N/A | N/A | N/A |
|  | 3F3832 | White | NR | N/A | N/A | N/A |
| **Close contacts** | 7768E4 | NR | NonHispanic | Day 2, Day 14, Day 28 | Negative, Negative, Negative | Negative, Negative, Negative |
|  | 61BBAD | White | NonHispanic | Day 0, Day 7, Day 14 | Negative, Negative, Negative | Negative, Negative, Negative |
|  | 80A16A | Asian | NonHispanic | Day 0, Day 14 | Negative, Negative | Negative, Negative |
|  | CE0CE8 | White | NonHispanic | Day 0, Day 7, Day 14 | Negative, Negative, Negative | Negative, Negative, Negative |
|  | 5ABDCB | White | NonHispanic | Day 0, Day 14, Day 28 | Negative, Negative, Negative | Negative, Negative, Negative |
|  | B96D3B | White | Hispanic | Day 0, Day 14, Day 28 | Negative, Negative, Negative | Negative, Negative, Negative |
|  | 6F894A | White | NonHispanic | Day 0, Day 7, Day 14 | Negative, Negative, Negative | Negative, Negative, Negative |
| **Mild** | 0B943B | White | NonHispanic | Day 0, Day 3, Day 14 | Negative, Negative, Positive | Positive, Positive, Positive |
|  | 2AD75E | Asian | NonHispanic | Day 0, Day 7, Day 14 | Negative, Negative, Borderline Positive | Negative<sup>&</sup>, Negative, Negative |
|  | 450905 | White | NonHispanic | Day 0, Day 7, Day 14 | Negative, Positive, Positive | Positive, Positive, Positive |
|  | BAAF62 | White | NonHispanic | Day 0, Day 7, Day 14 | Negative, Positive, Positive | Positive, Positive, Positive |
|  | 1A9B20 | White | NonHispanic | Day 0, Day 7, Day 14 | Negative, Positive, Positive | Positive, Positive, Positive |
|  | 75A2B6 | Asian | NonHispanic | Day 0, Day 7, Day 14 | Negative, Positive, Positive | Positive, Positive, Positive |
|  | DF309F | White | Hispanic | Day 0, Day 7, Day 14 | Negative, Positive, Positive | Negative<sup>&</sup>, Negative, Negative |
| **Pronounced** | 180E1A | White | NonHispanic | Day 0, Day 7, Day 14 | Negative, Positive, Positive | Positive, Negative, Negative |
|  | 82CCF5 | White | NonHispanic | Day 0, Day 7, Day 14 | Negative, Positive, Positive | Positive, Positive, Positive |
|  | B85D75 | White | NonHispanic | Day 0, Day 7 | Negative, Positive | Positive, Positive |
|  | 40067F | White | NonHispanic | Day 0, Day 14, Day 21 | Negative, Negative, Positive | Positive, Positive, Positive |
|  | 0BF51C | White | NonHispanic | Day 0, Day 7, Day 14 | Negative, Positive, NR | Positive, Positive, Positive |
|  | 0E1F8E | White | NonHispanic | Day 0, Day 7, Day 14 | Negative, Positive, Positive | Positive, Positive, Positive |
|  | 3F05F3 | White | NonHispanic | Day 0, Day 7, Day 28 | Negative, Positive, Positive | Positive, Positive, Negative |

\*NR = not reported
<sup>&</sup>Positive clinical PCR prior to enrollment, but negative research qPCR

**Supplemental Table 3.** Subject serology and PCR results on days when PBMCs were collected for single-cell assays. Serology testing was performed for IgG against the SARS-CoV-2 spike protein on each day of PBMC collection. Collection days are relative to study enrollment with sequencing profiling and specified for each subject.

| Disease Severity | Alias ID | Race | Ethnicity | Days of PBMC collection | Serology Test (IgG) | PCR |
| --- | --- | --- | --- | --- | --- | --- |
| **Healthy** | 22CBD6 | White | NR* | N/A | N/A | N/A |
|  | 77DA77 | White | NR | N/A | N/A | N/A |
|  | 4F2EAD | White | NR | N/A | N/A | N/A |
|  | F8FF1A | White | NR | N/A | N/A | N/A |
|  | 56E219 | Black | NR | N/A | N/A | N/A |
| **Close contacts** | 7768E4 | NR | NonHispanic | Day 2, Day 14, Day 28 | Negative, Negative, Negative | Negative, Negative, Negative |
|  | 61BBAD | White | NonHispanic | Day 0, Day 7, Day 14 | Negative, Negative, Negative | Negative, Negative, Negative |
|  | 80A16A | Asian | NonHispanic | Day 0, Day 14 | Negative, Negative | Negative, Negative |
| **Mild** | 0B943B | White | NonHispanic | Day 0, Day 3, Day 14 | Negative, Negative, Positive | Positive, Positive, Positive |
|  | 7085CA | White | NonHispanic | Day 0, Day 7, Day 14 | Negative, Negative, Positive | Negative<sup>&</sup>, Negative, Negative |
|  | 2AD75E | Asian | NonHispanic | Day 0, Day 7, Day 14 | Negative, Negative, Borderline Positive | Negative<sup>&</sup>, Negative, Negative |
|  | 450905 | White | NonHispanic | Day 0, Day 7, Day 14 | Negative, Positive, Positive | Positive, Positive, Positive |
|  | BAAF62 | White | NonHispanic | Day 0, Day 7, Day 14 | Negative, Positive, Positive | Positive, Positive, Positive |
| **Pronounced** | 180E1A | White | NonHispanic | Day 0, Day 7, Day 14 | Negative, Positive, Positive | Positive, Negative, Negative |
|  | 82CCF5 | White | NonHispanic | Day 0, Day 7, Day 14 | Negative, Positive, Positive | Positive, Positive, Positive |
|  | B85D75 | White | NonHispanic | Day 0, Day 7 | Negative, Positive | Positive, Positive |
|  | 40067F | White | NonHispanic | Day 0, Day 14, Day 21 | Negative, Negative, Positive | Positive, Positive, Positive |
|  | 0BF51C | White | NonHispanic | Day 0, Day 7, Day 14 | Negative, Positive, NR | Positive, Positive, Positive |

\*NR = not reported
<sup>&</sup>Positive clinical PCR prior to enrollment, but negative research qPCR

**Supplemental Table 4.** Independent cohort of subjects with critical symptoms requiring hospitalization. Summary of demographics for subjects with critical symptoms and associated clinical data including hospitalization, admittance to the ICU, and treatment with a respirator. PCR results were either from clinical or research testing, as noted.

| Disease Severity | Alias ID | Race | Ethnicity | Days of PBMC collection | Serology Test (IgG) | PCR | Hospitalized | ICU | Respirator |
| --- | --- | --- | --- | --- | --- | --- | --- | --- | --- |
| **Critical** | 45FBA5 | NR* | NR | Day 0 | Negative | Negative<sup>#</sup> | Y | Y | N |
|  | 2DB5F9 | Black-African American | NR | Day 0 | Positive | NR<sup>#+</sup> | Y | Y | Y |
|  | EB057C | Black-African American | NonHispanic | Day 0 | Positive | NR<sup>#+</sup> | Y | Y | Y |
|  | 936128 | White | NonHispanic | Day 0 | Positive | NR<sup>#+</sup> | Y | N | N |
|  | 460F8D | White | NonHispanic | Day 0 | Positive | Negative<sup>#</sup> | Y | N | N |
|  | 0993F8 | Black-African American | NonHispanic | Day 0 | Positive | NR<sup>#+</sup> | Y | N | N |

\*NR = not reported
<sup>#</sup>Treated in hospital and clinically adjudicated as COVID-19
<sup>+</sup>Positive clinical PCR, but no research qPCR reported

**Supplemental Figure 1.**
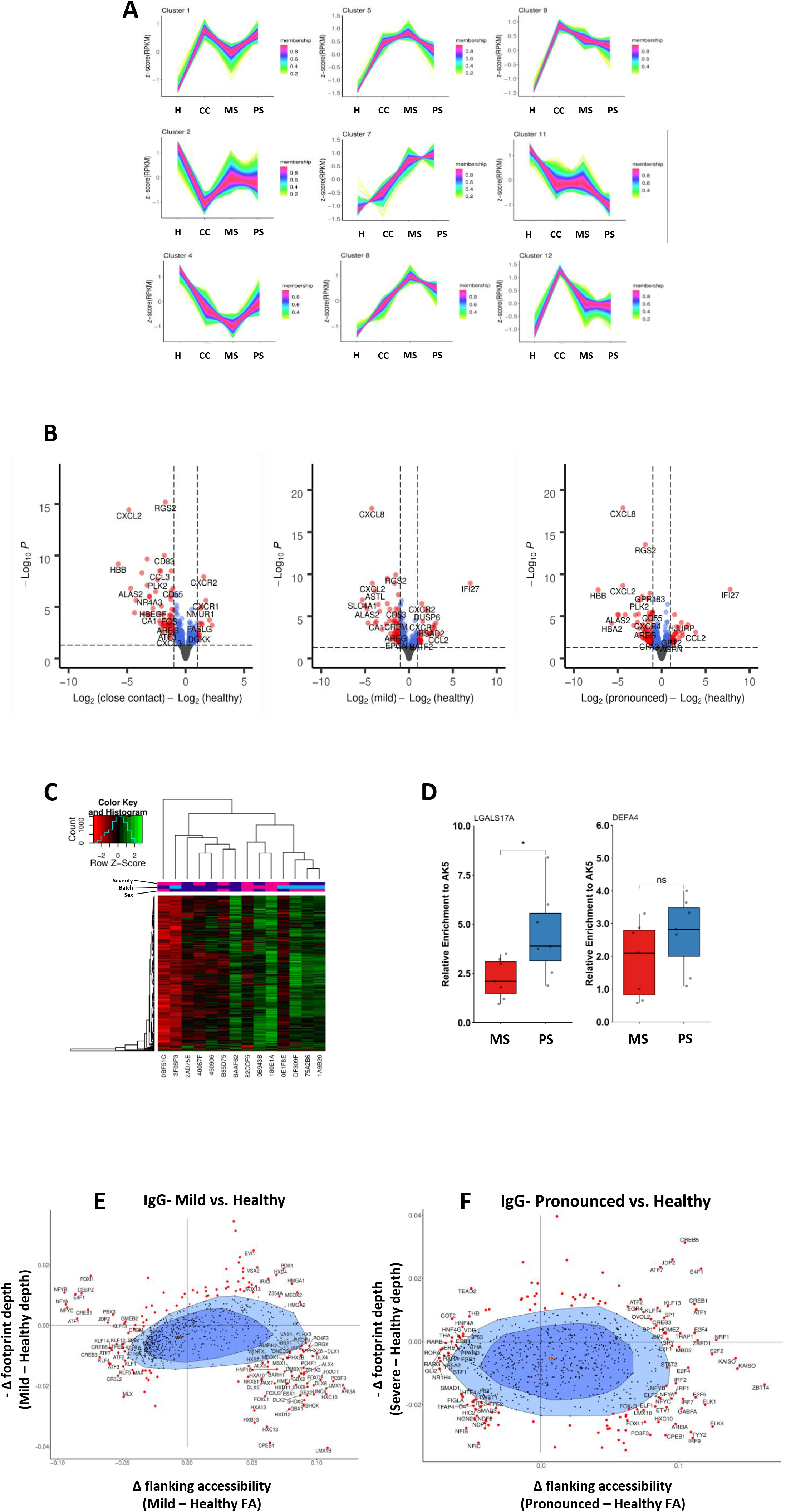
**(A)** Differentially accessible peaks clustered by fuzzy c-means clustering with n=12. Membership indicates similarity of a peak to the centroid of that cluster. **(B)** Differential gene expression from bulk RNA-seq comparing uninfected CC subjects vs. healthy controls (left), MS subjects vs. healthy controls (middle), and PS subjects vs. healthy controls (right). Only samples from the early (IgG-) timepoint were used. Significance is defined as adjusted p < 0.05 and absolute LFC > 1. **(C)** Heatmap of accessible chromatin peaks in seronegative MS and PS subjects. Significant peak (n=443) accessibility for subject. Severity annotation: purple = MS, pink = PS. Batch annotation: pink = batch 1, purple = batch 2, blue = batch 3. Sex annotation: pink = female, purple = male. **(D)** ATAC-PCR enrichment of PS-specific markers LGALS17A and DEFA4 relative to internal control AK5. ATAC-seq libraries from seronegative MS (n=7) and PS (n=7) subjects were used as input. *p < 0.05. **(E)** Bivariate genome footprinting analysis comparing transcription factor motif flanking accessibility and footprint depth in seronegative MS subjects to healthy controls. **(F)** Bivariate genome footprinting analysis comparing transcription factor motif flanking accessibility and footprint depth in seronegative PS subjects to healthy controls.

**Supplemental Figure 2.**
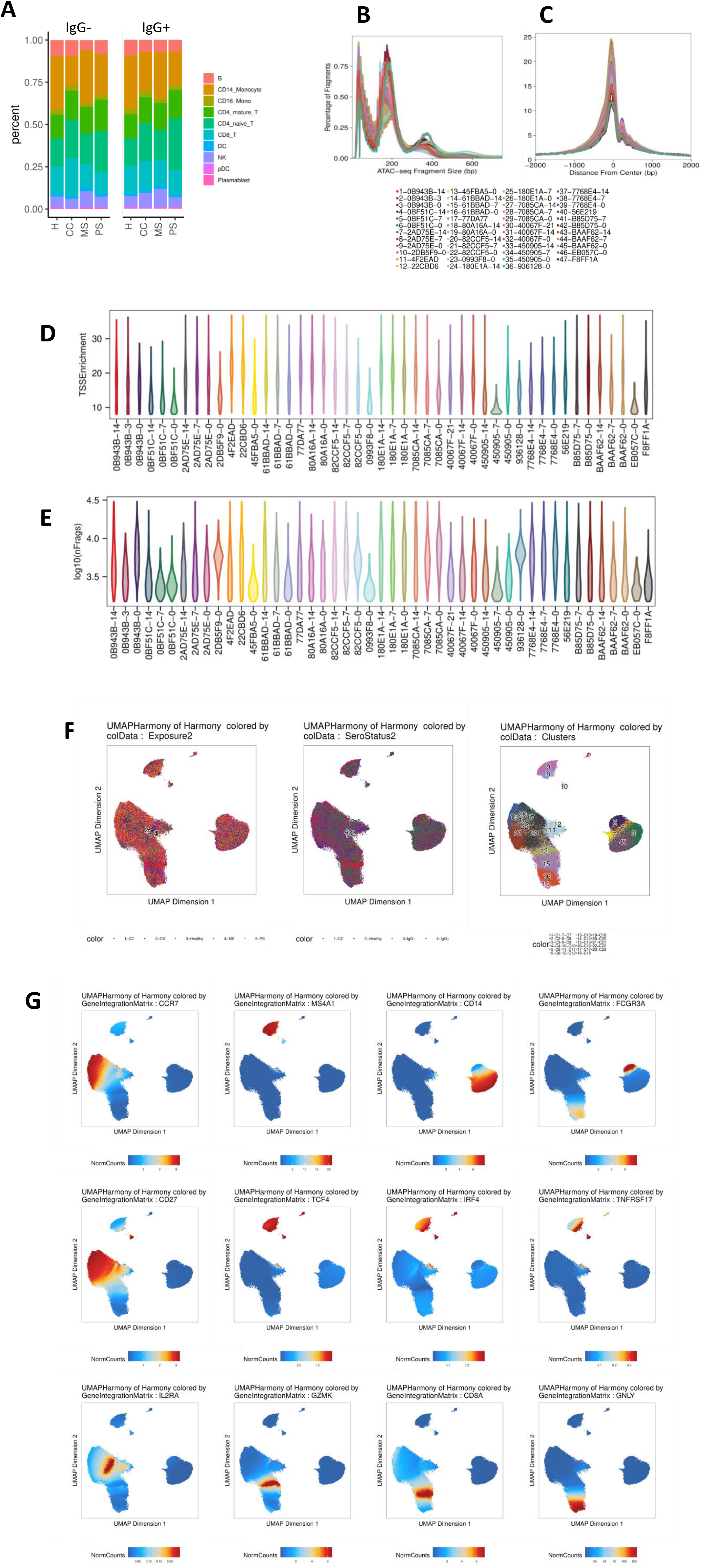
**(A)** Relative abundance of cell types represented in scATAC-seq cell atlas for each subject cohort split into IgG- and IgG+ timepoints. **(B)** ATAC-seq fragment size for each single-cell library. **(C)** Normalized insertion profile for each single-cell ATAC-seq library. **(D)** Distribution of TSS enrichment for cells in each single-cell ATAC-seq library. **(E)** Distribution of the log10(number of fragments) for cells in each single-cell ATAC-seq library. **(F)** UMAP representation of merged scATAC-seq cell atlas colored by subject cohort (left) and Seurat cluster (right). **(G)** UMAP heatmap of marker gene expression for major cell types represented in scATAC-seq after CCA label transfer from paired scRNA-seq datasets.

**Supplemental Figure 3.**
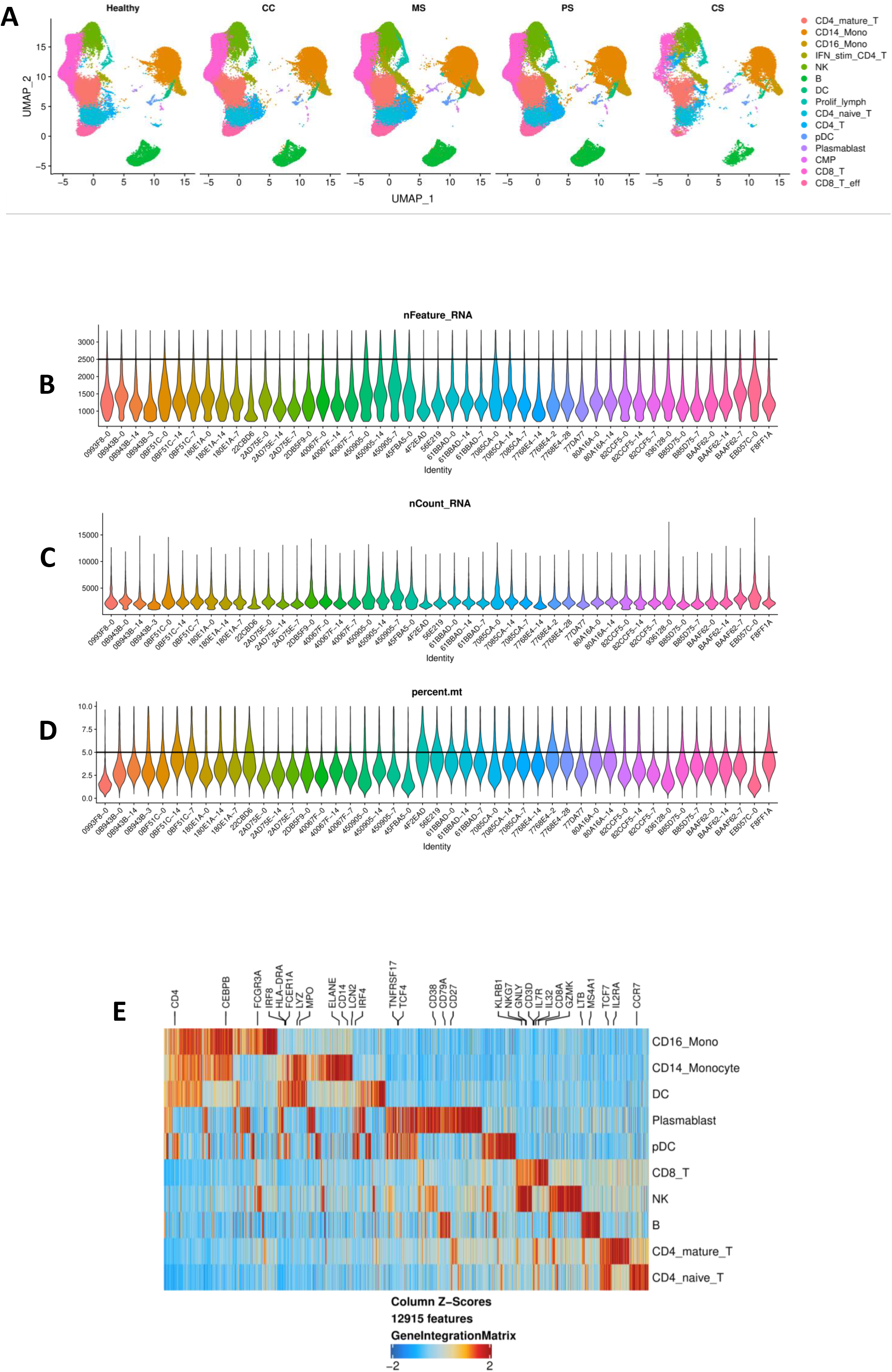
**(A)** Single-cell gene expression libraries prepared from each subject cohort. **(B)** Distribution of the number of genes represented in cells from each scRNA-seq library. Cells with > 2500 genes detected were removed from analysis. **(C)** Distribution of the number of unique molecular identifier (UMI) counts in cells from each scRNA-seq library. **(D)** Distribution of the percentage of mitochondrial reads in cells from each scRNA-seq library. Cells with > 5% of reads mapping to the mitochondrial genome were removed from analysis. (E) Cell type marker gene expression for profiled PBMCs. Column z-scores represent gene expression. Selected marker genes labeled for each cell type.

**Supplemental Figure 4.**
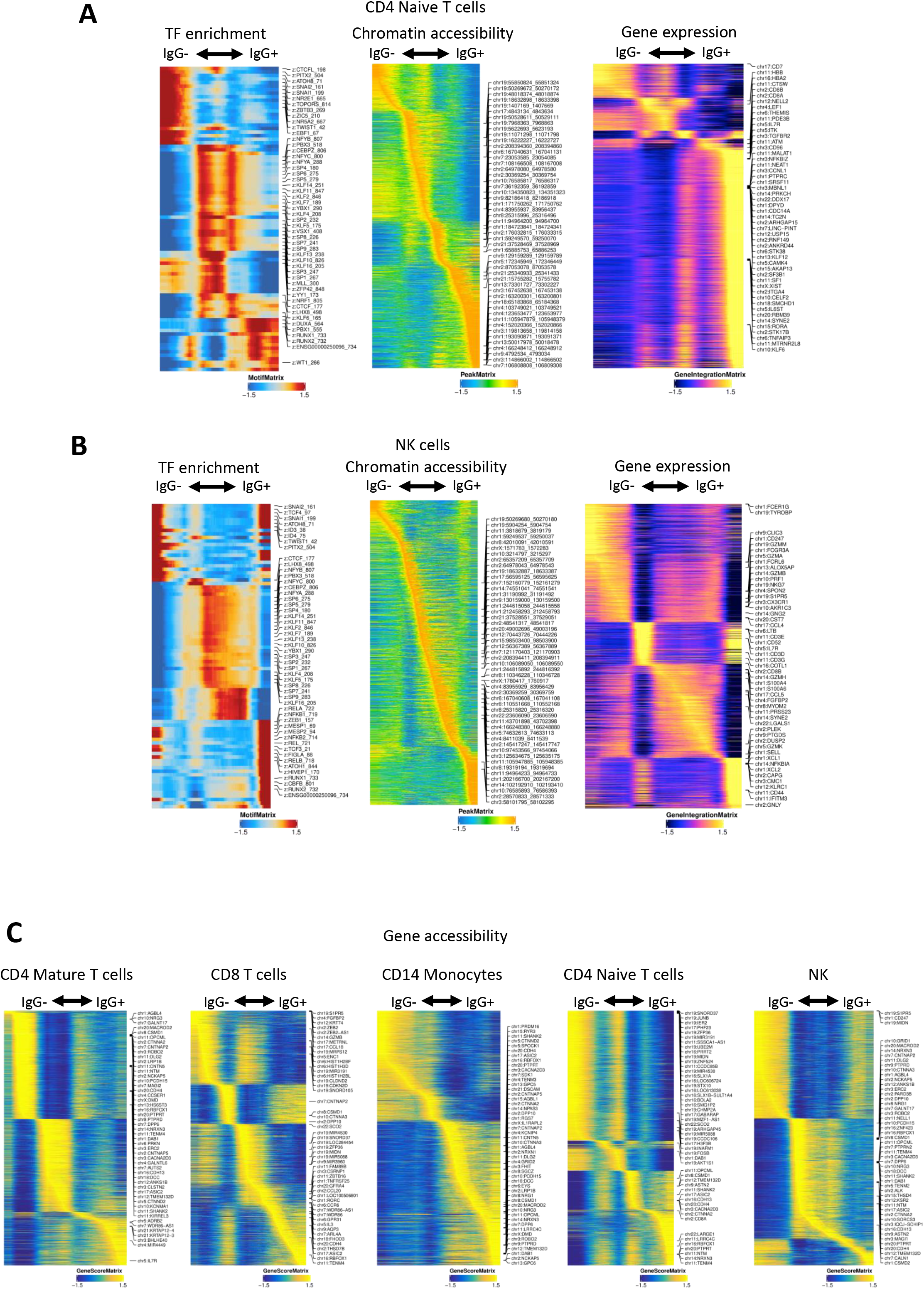
**(A-B)** Supervised trajectory analysis using CD4+ naive T cells **(A)** and natural killer (NK) cells **(B)** collected from both MS and PS subjects. Differential transcription factor motif enrichment, chromatin accessibility, and gene expression were correlated with seroconversion. **(C)** Differentially accessible genes identified using gene activity score estimation correlated with seroconversion in CD4+ mature T cells, CD8+ T cells, CD14+ monocytes, CD4+ naïve T cells, and NK cells (left to right).

**Supplemental Figure 5.**
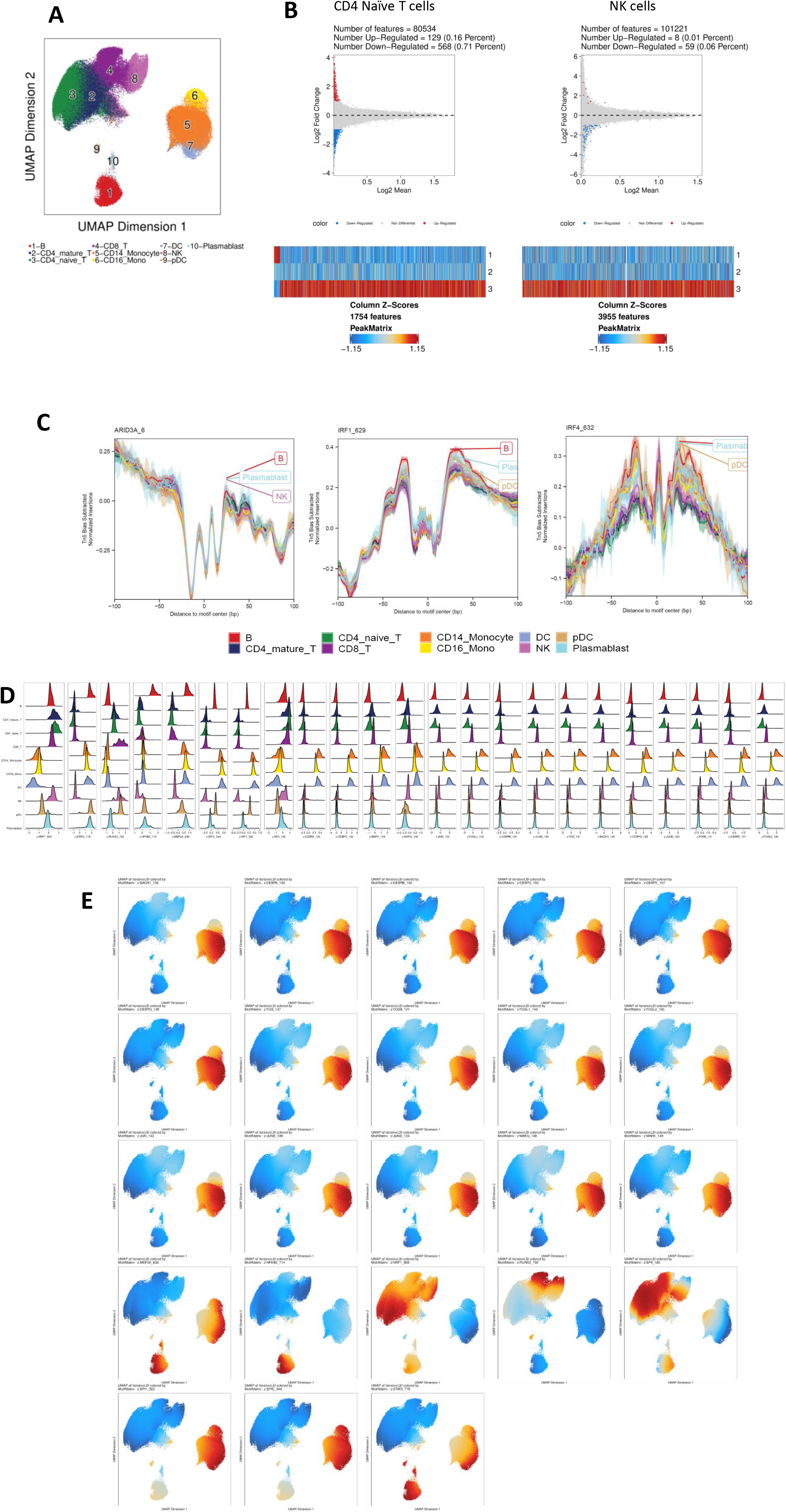
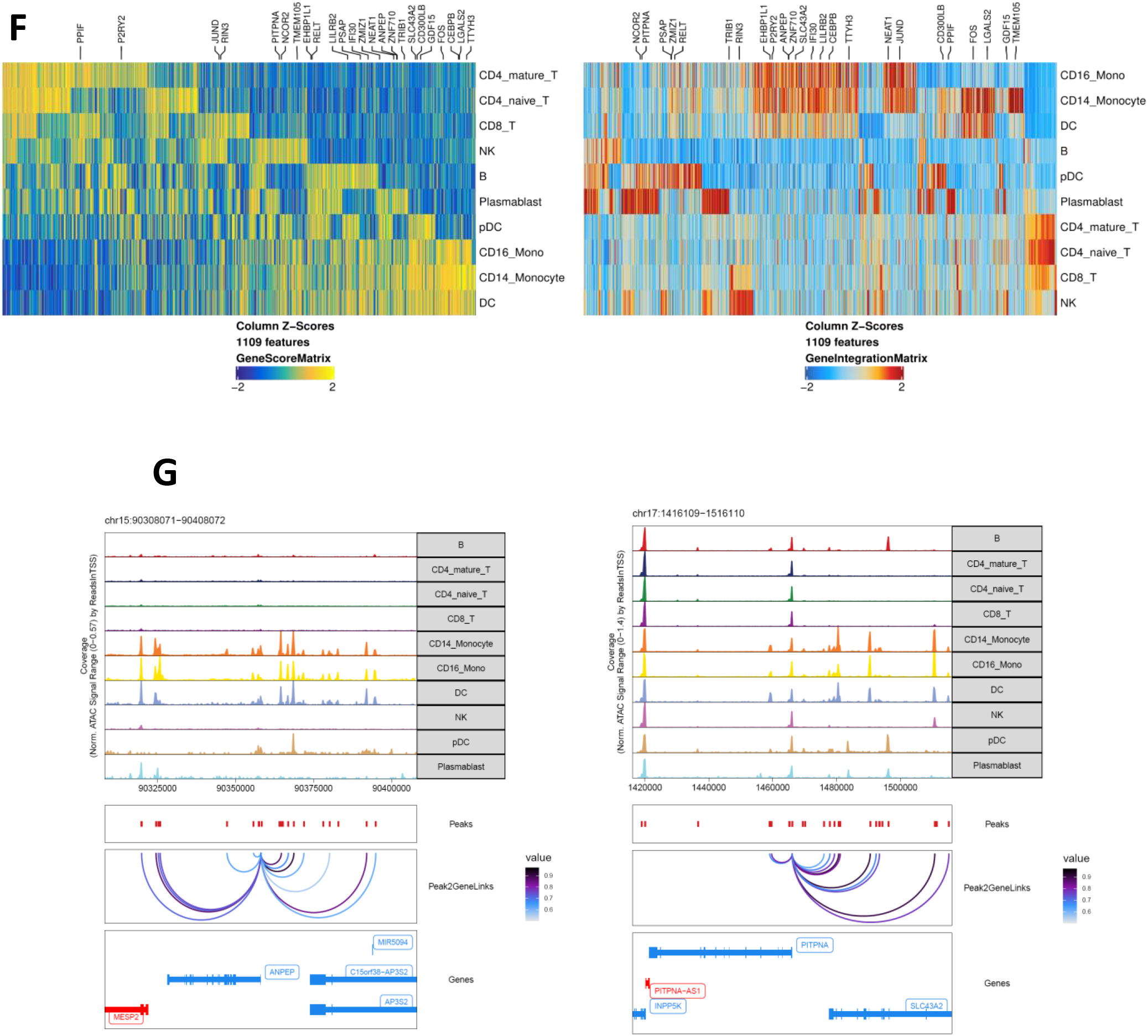
**(A)** UMAP representation of all cells collected from MS and PS subjects at early, mid, and late timepoints. **(B)** Differentially accessible peaksets identified in each major PBMC type at early vs. late timepoints (top MA plot) or associated with seroconversion (bottom heatmap). Left: CD4+ naïve T cells; Right: NK cells. **(C)** PS-specific transcription factor motif footprint occupancy was enriched in B cells, plasmablasts, NK cells, and pDCs. **(D)** Distribution of activator motif chromVAR accessibility deviation scores for each cell type collected from MS and PS subjects at all timepoints. **(E)** Activator motif accessibility heatmaps overlaid onto UMAPs of all cells collected from MS and PS subjects. **(F)** DORC gene activity (left) and gene expression (right) for 1109 loci plotted for each cell type. Labeled DORC genes are known to be regulated by a super-enhancer. **(G)** DORC genes regulated by super-enhancers have increased accessibility in CD14+ monocytes and other myeloid cells, including dendritic cells (DCs) and CD16+ monocytes. Peak-to-gene linkages are plotted with a correlation cutoff of 0.5.

**Supplemental Figure 6.**
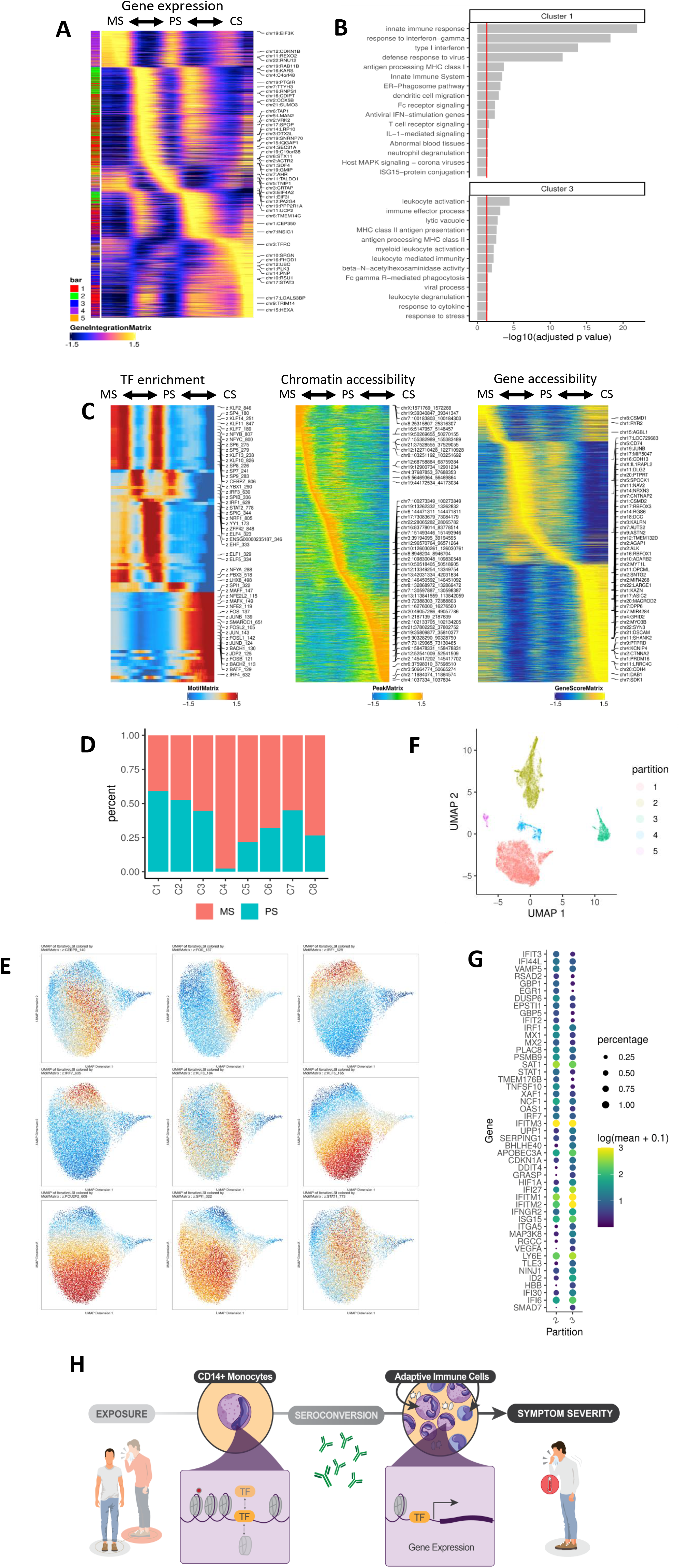
**(A)** Differential gene expression between CD14+ monocytes collected from seropositive MS, PS, and CS subjects. K-means clustering (k=5) plotted as row annotation. **(B)** Gene set enrichment analysis applied to clusters 1 and 3 which correspond to CD14+ monocytes from CS subjects. Red line indicates p = 0.05. **(C)** Supervised trajectory analysis using CD14+ monocytes collected from seropositive MS, PS, and CS subjects. Differential transcription factor motif enrichment, chromatin accessibility, and gene activity were associated with disease severity. **(D)** Percentage of cells in each scATAC-seq cluster from either MS or PS subjects. **(E)** Transcription factor motif accessibility of regulons identified using SCENIC represented as z-scores and overlaid on a UMAP representation of all CD14+ monocytes from MS and PS subjects. **(F)** scRNA-seq UMAP colored by Monocle 3 cell cluster partition. Partitions 2 (seronegative MS) and 3 (seronegative PS) are of interest. **(G)** Differential gene expression between partitions 2 and 3 plotted as percentage of cells expressing each gene and the log(mean + 0.1) gene expression. The top 25 markers for each partition are shown. **(H)** Schematic summary of a theory that CD14+ monocytes undergo chromatin remodeling prior to seroconversion, leading to downstream gene expression that impacts adaptive immunity and symptom severity.

